# Modeling post-logging height growth of black spruce forests by combining airborne LiDAR and historical forestry maps in eastern Canadian boreal forest

**DOI:** 10.1101/2021.01.22.427172

**Authors:** Batistin Bour, Victor Danneyrolles, Yan Boucher, Richard A. Fournier, Luc Guindon

## Abstract

Increase in forest disturbance due to land use as well as climate change has led to an expansion of young forests worldwide, which affects global carbon dynamics and forest management. In this study, we present a novel method that combines a single airborne LiDAR acquisition and historical harvesting maps to model height growth of post-logged black spruce-dominated forests in a 1700 km^2^ eastern Canadian boreal landscape. We developed a random forest model where forest height is a function of stand age, combined with environmental variables. Our results highlight the strong predictive power of this model: least-square regression between predicted and observed height of our validation dataset was very close to the 1:1 relation and strongly supported by validation metrics (*R*^*2*^ = 0.75; relative RMSE = 19%). Moreover, our findings indicated an ecological gradient responsible for differences in height growth at the landscape scale, with better growth rates on mesic slopes compared to badly drained soils on flat lands. With the increased availability of LiDAR data, this method is promising since it can be applied to forests across the globe that are affected by stand-replacing disturbances.

## Introduction

Over the last few decades, an increase in forest disturbance due to land use as well as climate change has led to the expansion of young forests worldwide (McDowell et al. 2020). This trend is likely to continue or even increase in the future (Boucher et al. 2017, McDowell et al. 2020). Thus, these young forests are playing an increasing and critical role in a variety of issues, for example, reaching a balance in global carbon dynamics (Cook-Patton et al. 2020) and maintaining forest ecosystem services (Kroll et al. 2020). These regenerating forests represent a critical stage in subsequent successional dynamics (Lindenmayer et al. 2019) and generally exhibit the highest growth rates. Yet, the dynamics of young forests have received surprisingly much less attention than have mature or old growth forests. More specifically, greater insights and better methods for modeling forest growth at such early stages of succession would considerably improve our ability to predict and manage changes in these forest landscapes.

Several factors may control young forest growth dynamics. For one, the time that has elapsed since the last stand-replacing disturbance (e.g., clearcutting, fire) plays an important role. Height growth rates are generally maximal in the early stages of succession, and tend to decline progressively with stand age as the trees attain their maximum height (Ryan et al. 2004). Yet, forest height growth is also mediated by a combination of environmental gradients operating at several scales. Regional climate plays an important role through three potential limiting factors: light, temperature and water (Boisvenue and Running 2006, Cook-Patton et al. 2020). In boreal forests, temperature is the main climatic limiting factor for growth with a short growing season (Huang et al. 2010), followed by regional drought events (D’Orangeville et al. 2018). Climatic gradients are also mediated by landscape-scale topographic gradients. For example, altitude, slope and exposure generate a diversity of local temperature characteristics that can influence the growth rates at the landscape scale (Nicklen et al. 2016). Similarly, site moisture conditions are strongly mediated by topography, surface deposits and drainage, with mesic mid- and upper-slopes generally leading to better tree growth rates when compared to poorly drained soils at lower slope positions (Lavoie et al. 2007, Laamrani et al. 2014).

There is an important and persistent tradition in ecology and forestry for the development of forest growth models (e.g., Vanclay and Skovsgaard 1997, Weiskittel et al. 2011, Pretzsch et al. 2015). Currently, most models are based on data gathered from extensive field measurements, such as long-term permanent plot networks (e.g., Pretzsch et al. 2014), or dendrochronological analyses of large numbers of trees (e.g., Huang et al. 2010, D’Orangeville et al. 2018). While acquisition of these data is generally time-consuming and expensive, the development of remote sensing methods to estimate forest structure characteristics offers cheaper alternatives, more specifically with respect to Light Detection And Ranging (LiDAR) (e.g., Næsset et al. 2013). Several studies have already proposed modeling forest growth using repeated airborne LiDAR acquisition (e.g., Meyer et al. 2013, Cao et al. 2016). Yet, these repeated acquisitions remain rather time-consuming and expensive since they imply a relevant time lapse between surveys (e.g., 5 to 10 years), which may further imply methodical challenges due to potential changes in LiDAR technological characteristics between surveys.

In this study, we present a simple novel approach that combines a single airborne LiDAR acquisition with historical harvesting maps to model forest height growth of post-logged boreal forests that are dominated by black spruce (*Picea mariana* [Mill.] BSP). Most sustainably managed forest landscapes include historical harvesting maps, particularly in even-aged managed stands (i.e., managed mostly through stand-replacing clearcuts; Lundmark et al. 2013). Our main objective was to develop and evaluate a predictive model of young forest stand height (10 to 50 years) as a function of stand age and other environmental explanatory variables.

## Materials and methods

### Study area

The study area covers 1,700 km^2^ in the black spruce-dominated, closed-crown boreal forests in the North Shore region of Quebec, eastern Canada (Fig. 1). Elevation ranges between 125 and 700 m and is associated with an important topographical gradient that includes lowlands and highland plateaus, and slopes that range between 0 and 20 degrees. The climate is typical of eastern Canadian boreal forest (Robitaille and Saucier 1998), with cold mean annual temperatures (−2.5 to 0°C) and abundant annual total precipitation (∼1300 mm). About two-thirds of our study area had been clearcut from 1955 to 2015 (Fig. 1). As is the case in most boreal forests, these stands are in remote areas that eventually are accessible only through very limited road networks a few years after harvesting because of rapid road degradation. Deterioration of the road network limits access, thereby making post-harvest field-based monitoring problematic. These characteristics make our study area a very good case study for developing and evaluating our new proposed growth modeling approach.

**Figure 1.**
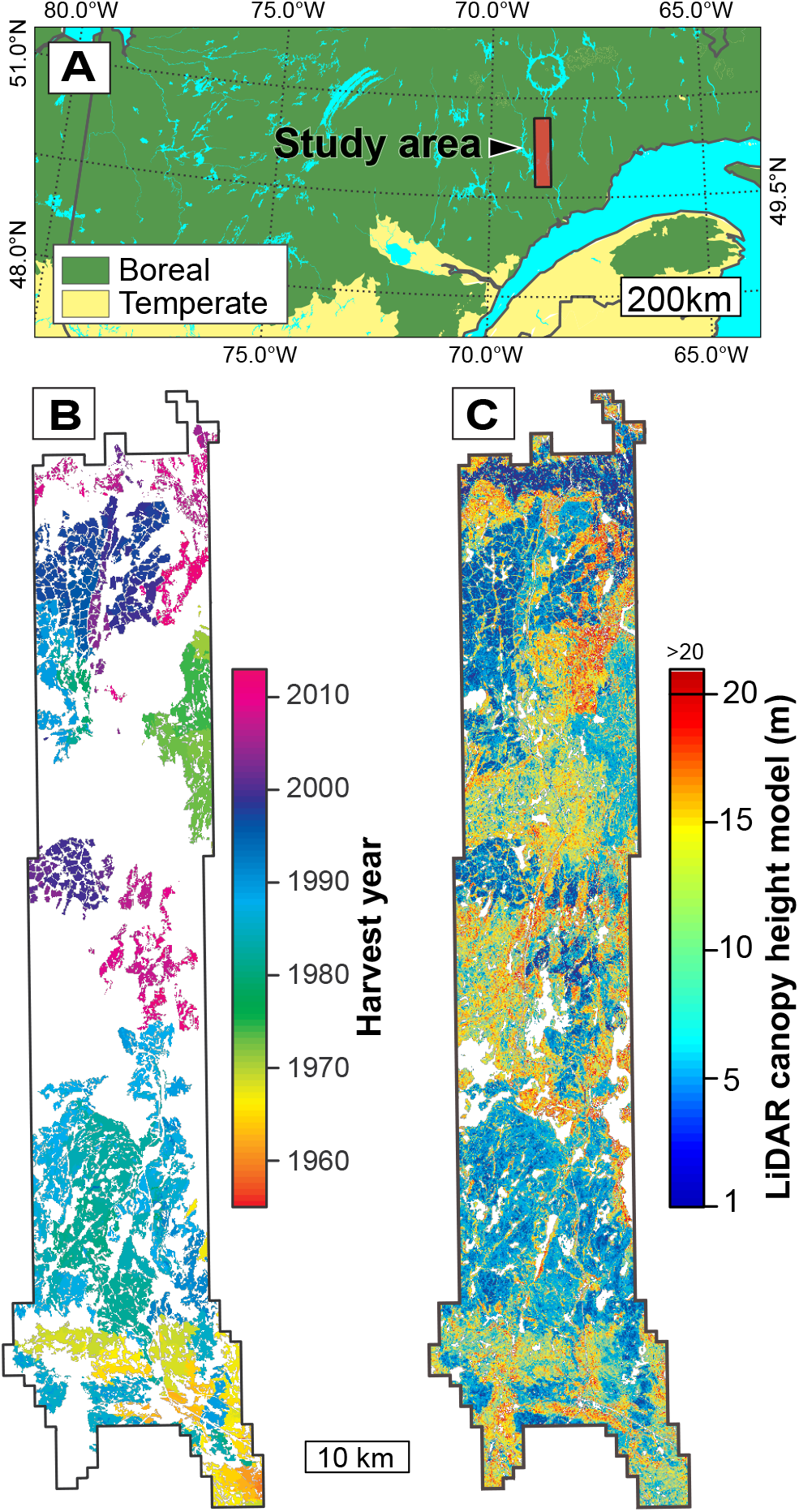
(A) Location of the study area in the boreal forest of eastern Canada. (B) a 20 m × 20 m raster layer of historical harvesting, and (C) the canopy height model based on airborne LiDAR data (2012 to 2016). Note that pixels > 20 m in (C) are displayed in dark red.

### Dataset description

The airborne LiDAR dataset was acquired from three campaigns (2012, 2014 and 2016) in which forests were overflown during or at the end of the growing season (June to November). About 77% of the study area had been surveyed in 2016, with the highest LiDAR point density (8.5 points.m^-2^), while the rest of the study area was sampled with lower point densities (6.6 points.m^-2^ in 2012 and 3.2 points.m^-2^ in 2014, which represented 18% and 5% of the study area, respectively). Further details on the LiDAR acquisition campaigns can be found in the Appendix S1 (Table S1).

Raw point clouds were first classified into ground and non-ground returns. A digital terrain model (DTM) was then fitted to the ground returns to produce a 20 m resolution raster. The DTM was subtracted from the elevations of all non-ground returns to produce a normalized point cloud. Finally, a canopy height model (CHM; Fig. 1) was obtained by using the 95^th^ percentile of point elevations of all non-ground returns (P95) in each 20 m × 20 m pixel, after removing returns < 1 m. P95 is frequently used to produce canopy height models (White et al. 2013), and exclusion of the lowest return (< 1 m) is usually applied to remove the returns from herbaceous-shrubby ground vegetation (Nyström et al. 2012).

The harvesting history (1955-2015) data were taken from forestry maps that are based on the interpretation of high resolution (1:20000 scale) aerial photographs and from annual harvesting reports (MFFP 2018). The information that was contained in the polygons was transformed into a 20 m × 20 m raster, matching the CHM data resolution (Fig. 1). The age of the trees within each logged pixel was then calculated as the difference between LiDAR acquisition year and harvesting year. Between 5 and 20 years following clearcutting, 24% of harvested stands were treated to precommercial thinning, a very common treatment in the boreal forest that reduces stand density and competing vegetation (Ashton and Kelty 2017). Consequently, we considered two distinct types of sylvicultural scenarios in our analysis: (1) clearcutting alone; and (2) clearcutting, followed by precommercial thinning.

Several additional environmental variables that could potentially influence forest height growth were also derived from LiDAR data or extracted from the forestry maps (Appendix S1: Table S2). Slope, exposure and a topographic wetness index (TWI; Beven and Kirkby 1979) were derived from the LiDAR DTM raster. Elevation, slope and exposure were then combined with historical meteorological data (1981-2010) to compute the mean growing degree-day (GDD) per 20 m × 20 m pixel with BIOSIM software (Régnière et al. 2014). Two categorical variables were extracted from modern forest maps: surface deposits (glacial, fluvio-glacial or rocky outcrops) and potential vegetation types. Potential vegetation types correspond to a fine scale level of Quebec’s forest classification system that refers to the late-successional vegetation that would be expected under given environmental conditions (climate, physiography; Grondin et al. 2014). In our study area, potential vegetation is represented by three major types: 1) balsam fir (*Abies balsamea* [L.] Miller)-black spruce forests (BF-BS); 2) balsam fir-paper birch (*Betula papyrifera* Marshall) forests (BF-PB), which are both found on rolling topography; and 3) black spruce-dominated forests on flat lands (BS).

We randomly sampled 20 m × 20 m pixels, where selected pixels must meet five conditions. First, because our analysis had focused on black spruce-dominated forests, only pixels with > 75% black spruce basal area, which was indicated in forest maps prior to clearcutting, were retained (MFFP 2018). Second, the first 50 m within the clearcut polygon boundaries were excluded to avoid border effects. Third, sampled pixels must be separated by a minimum distance of 250 m to avoid spatial autocorrelation (Matasci et al. 2018), the threshold of which was validated with Moran’s index. Fourth, only stands that were aged ≥ 10-years-old after clearcutting were retained, given that trees < 10-years-old could be confused with ericaceous shrubs, which can reach > 1 m in height (Matasci et al. 2018). Fifth, the 1:20000 polygons that identify clearcut areas had have a minimum size of 2 ha and could include small patches of remnant forest (< 0.05 ha). To remove these patches from the analysis, we excluded pixels with aberrant heights for a given age since they were very likely associated with remnant forest patches. Aberrant height thresholds were defined using a database of > 65,000 black spruce trees, the age and height of which have been measured in the field through Quebec’s network of permanent plots (MFFP 2016; Appendix S1: Fig. S1). The maximum height threshold for a given age was defined as the 95^th^ percentile of all field-based observations of tree height per age class. Applying these five conditions retained 3420 pixels that were subsequently allocated randomly to either a training set (2256 pixels; 66%) or a validation set (1164 pixels, 34%).

### Modeling forest height growth

Preliminary analysis involved identifying pairs of environmental explanatory variables that were ambiguously correlated. Problematic correlations (Pearson *r* > 0.5) were found between stand age, elevation and degree-days (Suppl. Fig. 2). This is not surprising since historically in this region, harvesting areas tended to progress over time from lower elevations in the southern part of our study area, to higher elevations in the northern part (Fig. 1, Suppl. Fig. 2). We decided to retain only stand age because it represented the most important gradient of values among these three variables for modeling forest height, which was confirmed by a generalized variance inflation factor analysis (Fox and Monette 1992; Appendix S1: Table S3).

We used a random forest model (Breiman 2001) to predict forest height growth since such machine learning approaches are very efficient in modeling non-linear ecological data with complex interactions (Christin et al. 2019). We trained the model using the *randomForest* function included in the *randomForest* package (version 4.6.14; Liaw and Wiener 2018) in the R statistical environment (R Core Team 2020). The training set (n = 2256) was analyzed to define optimal parameters using the *tuneRF* function, which was included in *randomForest* (Liaw and Wiener 2018). To evaluate the predictive power of our final model, we used our validation dataset (n = 1164) as a new input to the random forest model and compared observed and predicted values. We assessed the relative importance of variables in the model with the *importance* function of *randomForest*, which computes both the percentage increase in mean square error (%incMSE) and the increase in node purity for each explanatory variable (Liaw and Wiener 2018).

## Results

The comparison between observed and predicted pixel heights (i.e., LiDAR P95) from the validation dataset illustrates the strong predictive power of our random forest model (Fig. 2). The linear regression between predicted and observed values is very close to the theoretical relationship (1:1) and is strongly supported by several validation metrics (*R*^*2*^ = 0.74, relative root-mean-square-deviation = 18%, and mean error = 0.003 m).

**Figure 2.**
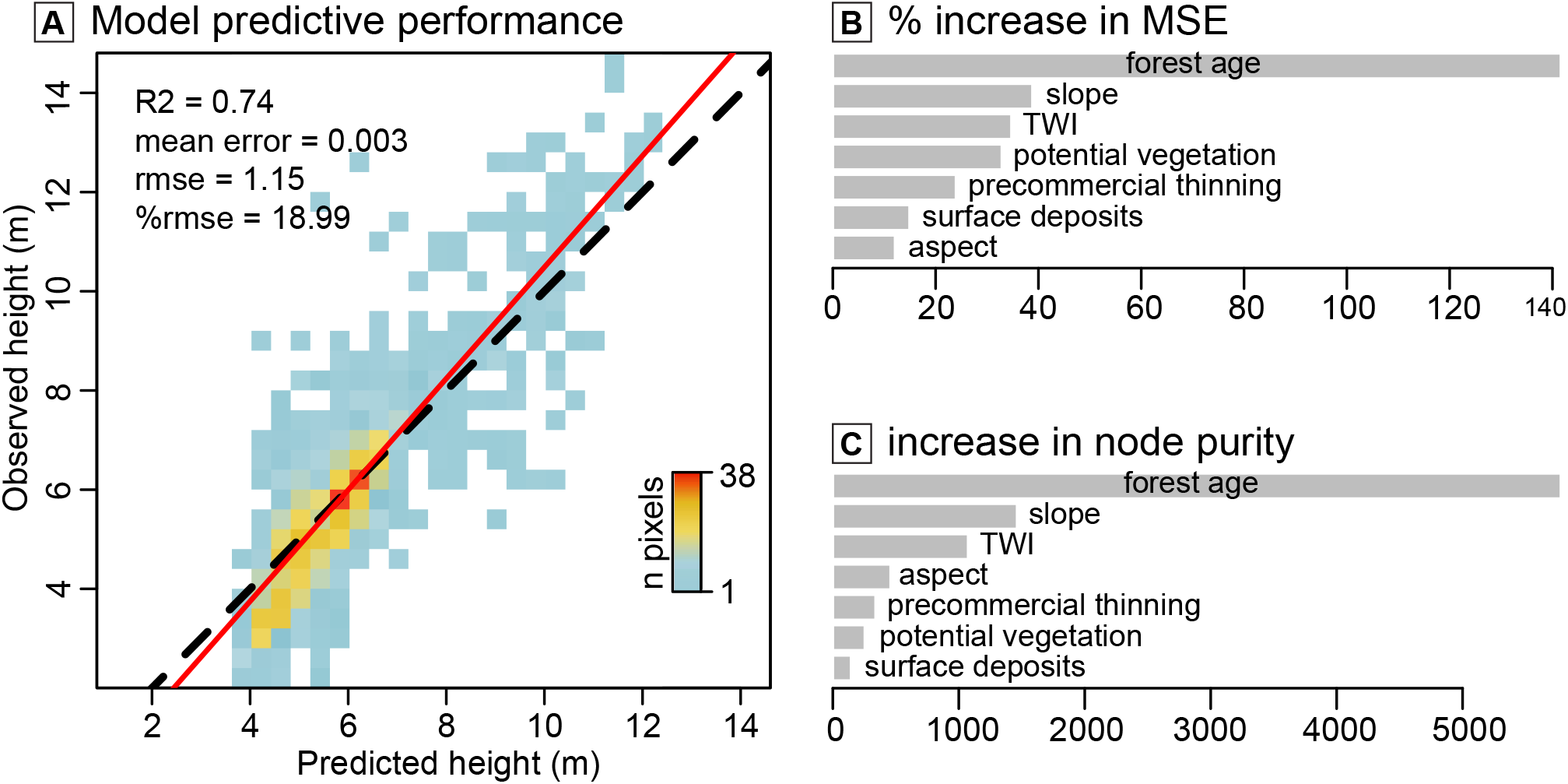
(A) Random forest model evaluation and (B, C) variable importance. Model predictive power was assessed in (A) through the comparison between observed and predicted pixel heights in the validation dataset (n = 1164 pixels). Point cloud density is displayed as a color gradient. The dotted black line shows the 1:1 theoretical relationship, while the solid red line shows the relationship modeled through ordinary least-squares regression. Variable importance in the random forest model was assessed (A) by percent increase in mean-square error and (B) by increase in node purity.

Application of the two tests (%incMSE and increase in node purity) within the random forest analysis leads to a similar ordering for the first three variables in terms of their relative importance and are relatively coherent for the other ones (Fig. 2). We have chosen to rank the relative importance of variables based on %incMSE, which is generally considered as the most reliable metric (Strobl et al. 2007). Stand age emerges as a dominant variable for predicting forest height (Fig. 2). Topographic characteristics emerge as secondary variables (slope and TWI; Fig. 2), with best height growth on slopes with high TWI (i.e., low moisture) compared to lower slopes with high TWI (i.e., high moisture; Fig 3). Potential vegetation types rank fourth (Fig. 2), with better growth on BF-PB sites (balsam fir-paper birch on rolling topography), compared to BF-BS and BS sites (balsam fir-black spruce forests on rolling topography and black spruce-dominated forests on flat lands, respectively; Fig. 3). The type of sylvicultural treatment ranks fifth (Fig. 2), with stands that have been treated to precommercial thinning showing slightly lower height growth compared to stands that have not been treated (Fig. 3). Surface deposits and aspect make the least important contributions in the model (Fig. 2); growth rates are generally higher on glacial surface deposits, while growth rates are generally lower on western and southeastern exposures (Fig. 3).

**Figure 3.**
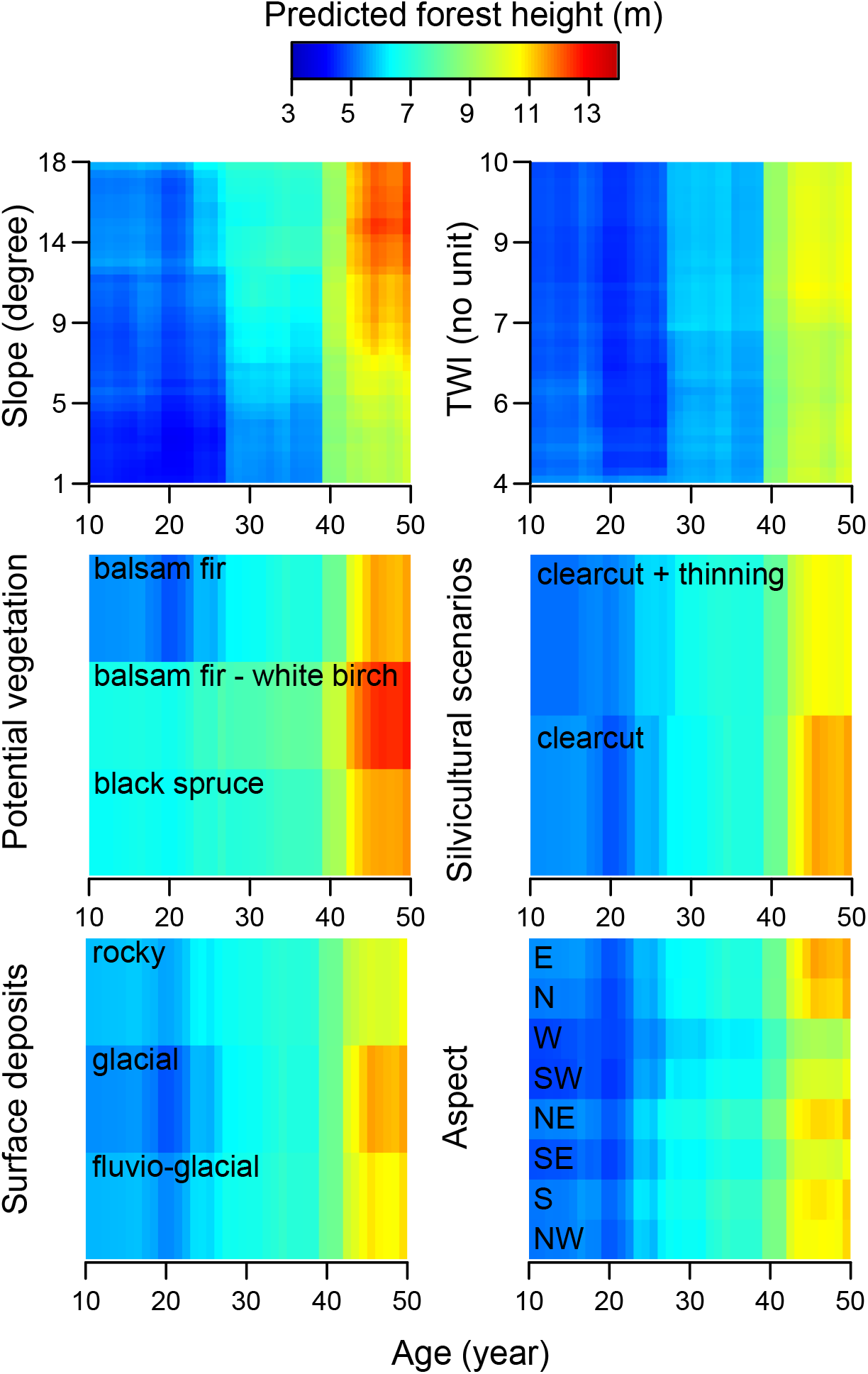
Interactive effects of stand age with other variables on forest height. For each variable, plots show the predicted forest height across the observed range of this variable, with other continuous variables held at median values (except for slope, which was held at 15° for categorical variable plots, to depict their effects in the best growing conditions). Categorical variables were held at the most common category across the training dataset (i.e., eastern exposure, clearcut silvicultural scenario, balsam fir potential vegetation, and glacial surface deposit). Ranges of slope and TWI were defined by their respective 2.5 and 97.5 percentiles.

## Discussion and conclusion

To our knowledge, this research presents the first study that combines airborne LiDAR data and historical disturbance maps to predict post-disturbance forest stand height growth. Overall, our results highlight the strong power of this approach: we were able to predict ≈75% of the validation dataset variation in stand height, with a mean error very close to zero. These highly supportive results strongly suggest that this approach could be generalized to larger areas and to other types of forests that are affected by stand-replacing disturbances (i.e., clearcuts, fire, hurricanes, agricultural land abandonment; Curtis et al. 2018). In our case study, the model highlighted an important ecological gradient that is responsible for differences in forest height growth at the landscape scale. Slope and site moisture (TWI) emerged as the second and third most important explanatory variables, after stand age. Best growth occurred on moderate slopes with low soil moisture compared to lower slopes with high soil moisture. This is not surprising since moist lower slopes are generally associated with poor drainage and high accumulations of organic matter that strongly limit forest productivity (Lavoie et al. 2007, Laamrani et al. 2014). Similarly, better growth rates were found on balsam fir-paper birch potential vegetation types (BF-PB) and glacial surface deposits that are likely associated with this drainage and organic matter gradient, given that these sites are generally associated with best drainage conditions.

Our results also highlighted the potential of our method to model the effects of different stand-replacing disturbance types on forest height growth. Precommercial thinning after clearcutting had a small but significant negative effect on height growth when compared to other stands, which could be linked to several mechanisms. First, although we made efforts to limit our analyses to black spruce-dominated stands (> 75% of the basal area), the presence of a minor deciduous component is ubiquitous in our data (Suppl. Fig. 3). The slower growth rates that were observed in precommercial thinning scenarios may be partly linked to the goal of precommercial thinning, which removes fast-growing deciduous species. These thinned individuals include mostly *Betula papyrifera*, and to a lesser extent, trembling aspen (*Populus tremuloides* Michaux). Indeed, a lower proportion of deciduous components are encountered in thinned stands (Suppl. Fig. 3). Slower growth rates in thinned stands could also be explained by a short-term negative effect that is exerted by thinning on forest height growth, followed by positive effects over much long-term periods (>30 years after thinning; Zhang et al. 2006) that are not perceived in our analysis.

As stand-replacing disturbance increases due to land-use and climate change worldwide, young stands are playing an increasing and critical role in global carbon dynamics and forest management (Cook-Patton et al. 2020, McDowell et al. 2020). There is a pressing need for greater knowledge of such early successional forest stages, which generally exhibit higher growth rates compared to mature or old growth forests. Our method combines airborne LiDAR and historical stand-replacing disturbance maps and, thus, provides a very simple and powerful approach to modeling young forest growth. This method appears particularly promising, given that many countries are currently building extensive national LiDAR inventories, since the technique could be applied at larger scale and virtually to any forest worldwide that is affected by stand-replacing disturbances (e.g., clearcuts, fire, hurricanes, agricultural land abandonment; Curtis et al. 2018). Such data are even becoming available at the global scale with space-borne LiDAR forest structure and aboveground biomass data (Hancock et al. 2019), together with remote-sensed historical forest disturbance areas (Hansen et al. 2013) and types (Guindon et al. 2017, 2018, Curtis et al. 2018). Applying our method to larger areas would allow the construction of improved models that should have broad-scale applications in ecology and forestry. For example, this effort could permit modeling effects of large-scale climate gradients and help to predict how changing climate would alter future forest growth.

## Acknowledgements

This project was financially supported by the Collaborative Research and Development program of the Natural Sciences and Engineering Research Council (NSERC grant # RDCPJ 508853-17), through a grant entitled “Outils spatiaux de gestion des forêts boréales après-feu et d’accès aux territoires dans un contexte de changements globaux: régénération après-feu.” We are grateful to our industrial partners: Ryam, Produits forestiers résolu, Rébec, Barrette-Chapais, and Rival solutions. This research could not have been performed without the collaboration of the Quebec Ministère des Forêts de la Faune et des Parcs (MFFP), and more specifically, J.F. Bourdon for the availability of high quality datasets (forest plots and LiDAR) and his support of the data analysis. Also, we wish to thank J. Noël and I. Auger of the MFFP for their support related to the statistical analysis, P. Villemaire of Natural Resources Canada for the availability and analysis of several explanatory variables and W.F.J. Parsons for the English revision.

