## Appendix 1 for "Modeling post-logging height growth of black spruce forests by combining airborne LiDAR and historical forestry maps in eastern Canadian boreal forest"

**Appendix S1**

**Table S1.** LiDAR survey details.

| Year of acquisition | 2012 | 2014 | 2016 |
|---|---|---|---|
| Period of the year | July to August | June to July | July to November |
| LiDAR technology | Riegl LMS-Q680i | Riegl LMS-Q680i | Optech ALTM Galaxy |
| Impulse frequency | 160 Hz | 87 Hz | 350 Hz |
| Scanning frequency | 108 Hz | 69 Hz | 52 Hz |
| Flight altitude | 600 m | 800 m | 1200 m |
| Flight speed | 51 $m.s^{-1}$ | 51 $m.s^{-1}$ | 72 $m.s^{-1}$ |
| Wipe angle | 60° | 60° | 24° |
| Mean point density | 6.6 $points.m^{-2}$ | 3.2 $points.m^{-2}$ | 8.5 $points.m^{-2}$ |
| % of study area | 18% | 5% | 77% |

**Table S2.** Description of variable sources, type (Cont., continuous; Categ., categorical), and range. In the range column of continuous variables, the first number is the mean value, while numbers within parentheses are minimum and maximum values.

| **Variables** | **Source** | **Type (unit)** | **Range** |
|---|---|---|---|
| Stand height | LiDAR | Cont. (m) | 6.04 (1.79 - 16.19) |
| Stand age | Forest maps | Cont. (year) | 29.15 (3 - 53) |
| Elevation | LiDAR | Cont. (m a.s.l.) | 460 (130 - 700) |
| Slope | LiDAR | Cont. (°) | 7.82 (0.01 - 28.49) |
| TWI | LiDAR | Cont. (no unit) | 6.26 (3.31 - 14.79) |
| Exposition | LiDAR | Categ. | N, NE, E, SE, S, SO, O, NO |
| Degree-days | Meteorological | Cont. (°C) | 1123 (990 - 1289) |
| Sylvicultural scenarios | Forest maps | Categ. | Clearcut, Clearcut + thinning |
| Potential vegetation | Forest maps | Categ. | BF-BS, BF-PB, BS |
| Surface deposit | Forest maps | Categ. | Glacial, fluvio-glacial, rocky |

**Table S3.** Generalized variance inflation factor (GVIF) analysis. GVIF analysis (Fox and Monette 1992) is analogous to classical variance inflation factor (VIF) analysis, but allows calculation of GVIF for categorical variables, where values can be compared with values of continuous variables after correcting for differences in degrees-of-freedom (Df). The transformation that permits comparison of categorical and continuous variables is $GVIF^{(1/(2\times Df))}$. To apply a classical threshold of VIF <10, $GVIF^{(1/(2\times Df))}$ should be less than $10^{(1/(2\times Df))}$. The selection column indicates variable rejection after applying a threshold equivalent to VIF > 10, while all displayed values for accepted variables are smaller than an equivalent VIF threshold of 5.

| | GVIF | Df | $GVIF^{(1/(2\times Df))}$ | Selection |
|---|---|---|---|---|
| Age | 1.9615 | 1 | 1.4005 | YES |
| Potential vegetation | 1.5855 | 2 | 1.1221 | YES |
| Surface deposits | 1.6953 | 2 | 1.1411 | YES |
| Sylvicultural scenarios | 1.1617 | 1 | 1.0778 | YES |
| Slope | 1.3901 | 1 | 1.1790 | YES |
| Degree-days | 13.3901 | 1 | 3.6592 | NO |
| TWI | 1.3077 | 1 | 1.1435 | YES |
| Elevation | 11.6518 | 1 | 3.4135 | NO |
| Aspect | 1.0553 | 7 | 1.0039 | YES |

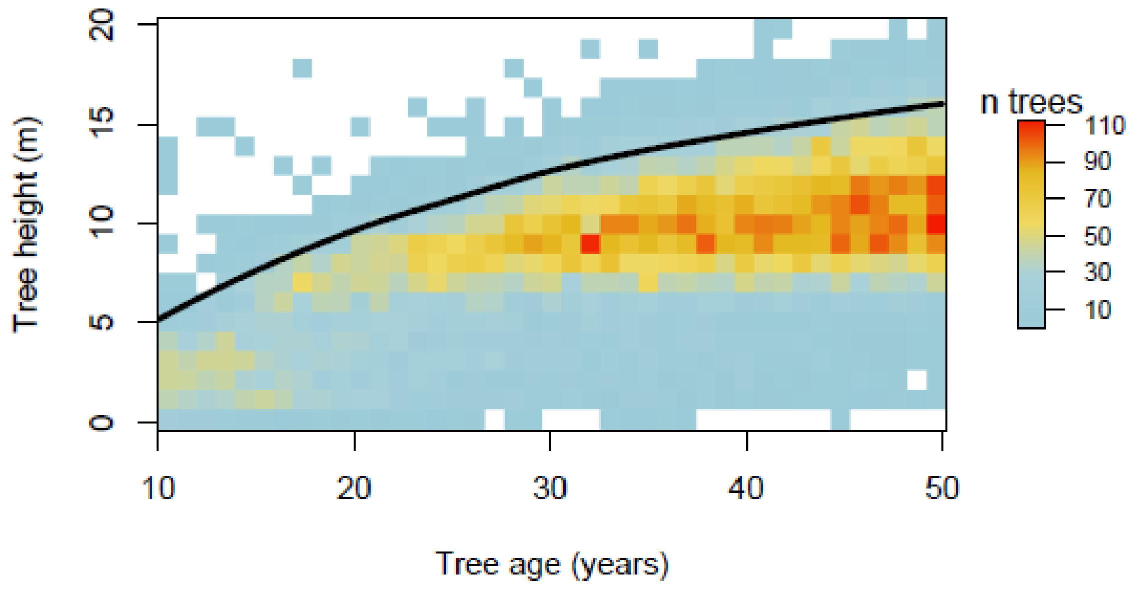


**Figure S1.** Age-height relationship established for 65000 black spruce trees that were measured as part of the permanent plot inventory network maintained by the provincial government of Quebec. The black line shows the 95th percentile that marked an aberrant height threshold for each age class.

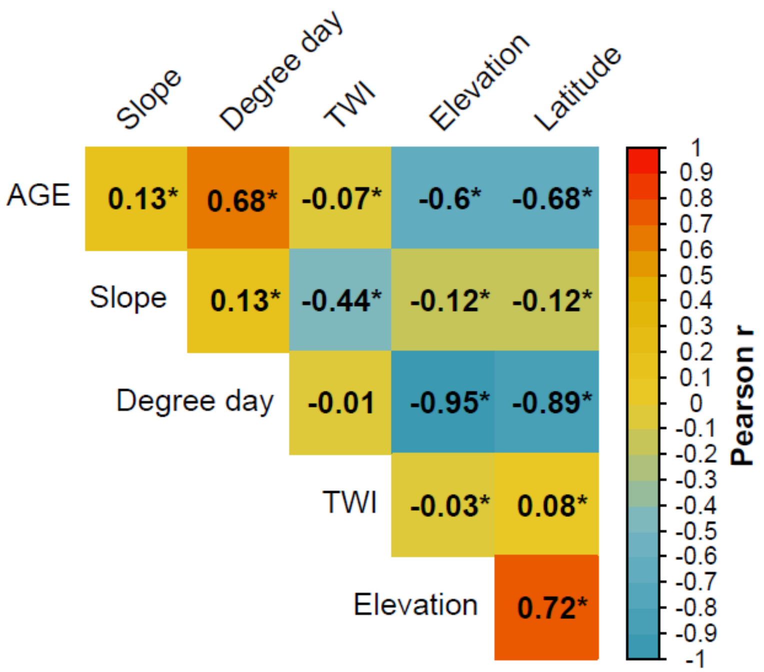


**Figure S2.** Correlation matrix between continuous and categorical variables; the value of Pearson's r is specified for each pair of variables. Latitude has been added to the list of variables to obtain information on the effect of the south-north gradient of the study site. The cell color depicts strength of correlation; (*) indicates relationships where $p < 0.05$.

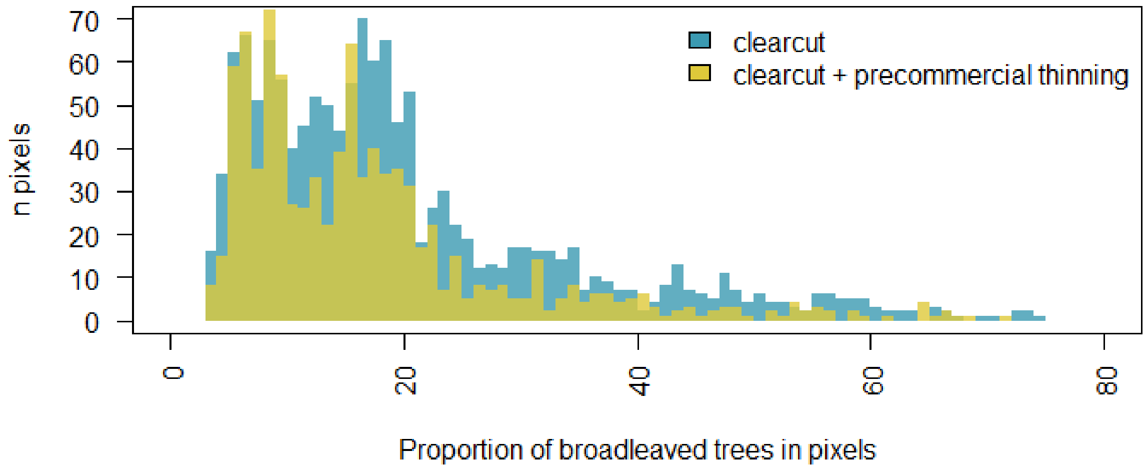


**Figure S3.** Proportion of deciduous trees in the pixels of the training dataset. The proportion of deciduous trees was established by the Laurentian Forestry Centre (Canadian Forest Service) by a random Forest classification of the territory between the provinces of Ontario and Newfoundland and Labrador (30 m × 30 m raster).
